# Profiling gene expression of the host response to a *Plasmodium vivax* irradiated sporozoite immunization and infectious challenge

**DOI:** 10.1101/501957

**Authors:** Monica L. Rojas-Peña, Dalia Arafat, Juan Manuel Velasquez, Swetha Garimalla, Myriam Arévalo-Herrera, Sócrates Herrera, Greg Gibson

## Abstract

The development of vaccines that provide sterile protection against human malaria is a major global public health priority requiring a better understanding of the mechanisms involved in natural and vaccine-induced sterile immunity. RNAseq was used to profile gene expression of peripheral blood samples from 12 Duffy positive (Fy+) (*Plasmodium vivax* susceptible) volunteers enrolled in a phase 2 trial who were vaccinated with radiation attenuated *P. vivax* sporozoites (RAS; 5 were protected, 7 not) and from 5 Fy- (*P. vivax* resistant) volunteers exposed to mosquitoes harboring live non-attenuated sporozoites. Blood samples were obtained before and after immunization as well as after controlled infection with live *P vivax* sporozoites. The most profound changes in gene expression were observed between baseline and post-challenge, with 97 distinct signatures differentiating protected and not protected Fy+ individuals. Differentiation was also observed between Fy- and Fy+ protected individuals, notably with downregulation of multiple inflammatory responses as well as extracellular matrix-related gene activity. Analysis of transcriptional modules shows that both B-cell and T-cell signaling are reduced while cell cycle regulation, interferon response, and other informative signatures are elevated in individuals who are not protected against malaria. An asymptomatic individual had an intermediate profile indicative of differential transcription associated with pathology and symptomology. Systems biology thus provides insight into how whole malaria-attenuated sporozoites prime the immune system to protect against malaria, as well as the transcription responses that are associated with sterile protection.

**AUTHOR SUMMARY:** Transcriptomic analysis has shed considerable light on the molecular mechanisms of vaccine efficacy against a variety of viral pathogens, but has not yet been used in relation to parasite vaccinology. Here we describe whole peripheral blood gene expression profiling to contrast the contributions of myeloid and lymphoid immune responses to vaccine-induced and natural protection against *Plasmodium vivax-* mediated malaria. Two modes of modular bioinformatic analyses are applied to RNAseq data from pre- and post-vaccination samples from 19 volunteer participants in a clinical trial investigating the efficacy of an irradiated sporozoite vaccine. During active infection, both B- and T-cell activity are reduced in individuals who are not protected by vaccination relative to those who are protected. Lack of the Fy antigen prevents entry of parasite into red blood cells, thereby providing natural clinical immunity, but we also document differences in T-cell and neutrophil activity in these individuals relative to those protected by irradiation of the parasite. One individual who was asymptomatic for malaria but had blood parasite despite vaccination showed an intermediate response indicating how systems biology approaches can distinguish gene expression associated with pathology and infection.

## INTRODUCTION

Development of effective vaccines remains an important component of the strategy to decrease and potentially eliminate malaria [1,2]. Human malaria is caused by at least five different parasite species of the genus *Plasmodium*, which display distinct biology and disease manifestations. *P. falciparum* and *P. vivax* are the most relevant to public health worldwide [3], the latter accounting for ~20% of the global malaria burden and producing a chronic disease that may develop to complicated and severe cases with infrequent lethality [4,5].

Multiple genetic traits partially protect humans from severe clinical manifestations or completely prevent blood parasite development [6], thereby reducing disease in infected individuals [7]. For example, lack of expression of the Duffy (Fy) blood group in African populations associates with protection from *P. vivax* blood infection [8,9] and consequently from the clinical manifestations associated with this parasite stage, despite its full development in the liver. This species also has the capacity to remain dormant as hypnozoites in the liver and periodically reactivate, inducing clinical relapses which represent an enormous challenge for *P. vivax* control and elimination and highlights the importance of developing vaccines targeting the parasite liver forms. Rational design of malaria vaccines is likely to be accelerated by enhanced understanding of the molecular mechanisms involved in the clinical immunity developed in individuals permanently exposed to infection in endemic areas [10,11], or by sterile immunity experimentally induced by vaccination with attenuated parasites [12].

In recent years, we have developed a controlled human malaria infection model with *P. vivax* sporozoites (CHMI) in Colombia [13–15], which supported gene expression profiling of the early response to *P. vivax* in malaria-naïve and semi-immune volunteers [16]. More recently, a 20-volunteer Phase I/II clinical trial of a *P. vivax* radiation-attenuated sporozoite (RAS) vaccine assessed the protective efficacy against *P. vivax* sporozoite infectious challenge [17].

Systems biology approaches are being implemented to enhance the knowledge and understanding of immunity acquisition to infectious agents [18,19], and gene expression profiling has been used to gain insight into natural clinical protection to viral diseases [17,20,21], as well as for identifying gene signatures associated with vaccination response [22–25]. However, while it is clear that multiple arms of the immune system are engaged in this protection, the precise mechanisms leading to immunity remain to be elucidated. In malaria, the pre-erythrocytic liver parasite stage, is thought to be an optimal vaccine target because during this phase, the infection remains clinically silent; once the parasite is released to the blood stream (erythrocytic stage) malaria clinical manifestations develop. Gene expression profiling of leukocytes from humans rendered immunologically sterile by RAS experimental vaccination is likely to provide valuable insight for rational subunit vaccine development.

This study focuses on transcriptomic profiles of human leukocytes to establish the nature of differences in the immune profiles between: (i) Fy+ protected and non-protected by RAS vaccination, and (ii) experimentally and naturally (Fy-) refractory individuals exposed to immunization with non-attenuated sporozoites. Knowledge of the co-regulation of gene expression in blood is used to partition the transcriptional profiles into contributions of various myeloid and lymphoid cell-types.

## RESULTS

Peripheral blood gene expression profiles were obtained for 20 volunteers who completed a single blind randomized clinical trial of RAS immunization [17], as schematized in Fig 1. Five individuals were Fy-, and 15 were Fy+ (12 RAS immunized and three mock-vaccination controls); of the RAS group, 5 attained sterile immunity, 6 contracted malaria, and 1 was infected but asymptomatic. Table 1 shows the demographics and clinical attributes of these volunteers. The final RNAseq dataset includes Baseline, Pre-challenge and Diagnosis day time points with exclusion of five samples as reported in S1 Table. It consists of 55 RNAseq single end 100bp samples with a total of 13,282 genes that had at least one cpm in at least 3 of the samples.

**Figure 1.**
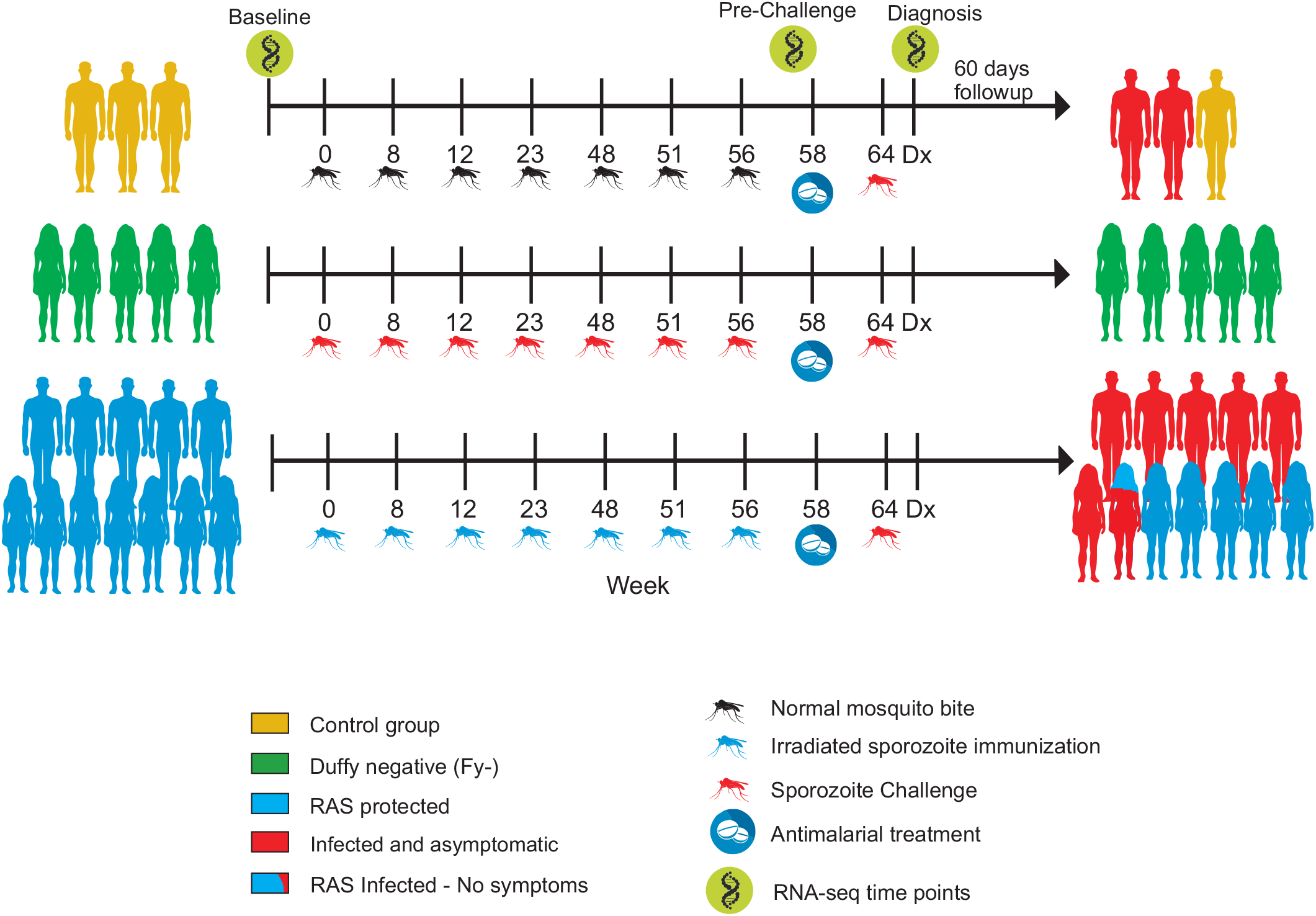
Experimental design and sample collection timeline. 58 total samples were used for RNAseq, taken at three time points represented by the green symbols: Baseline (20 samples), Prechallenge (19 Samples) and day of Diagnosis (19 samples). Controls, Fy^-^ and Fy^+^ Vaccinated individuals are represented by yellow, green and blue silhouettes of representative genders on the left, and red shading on the right indicates which individuals were diagnosed with malaria. One woman (red/blue) was positive for infection but asymptomatic.

**Table 1.**
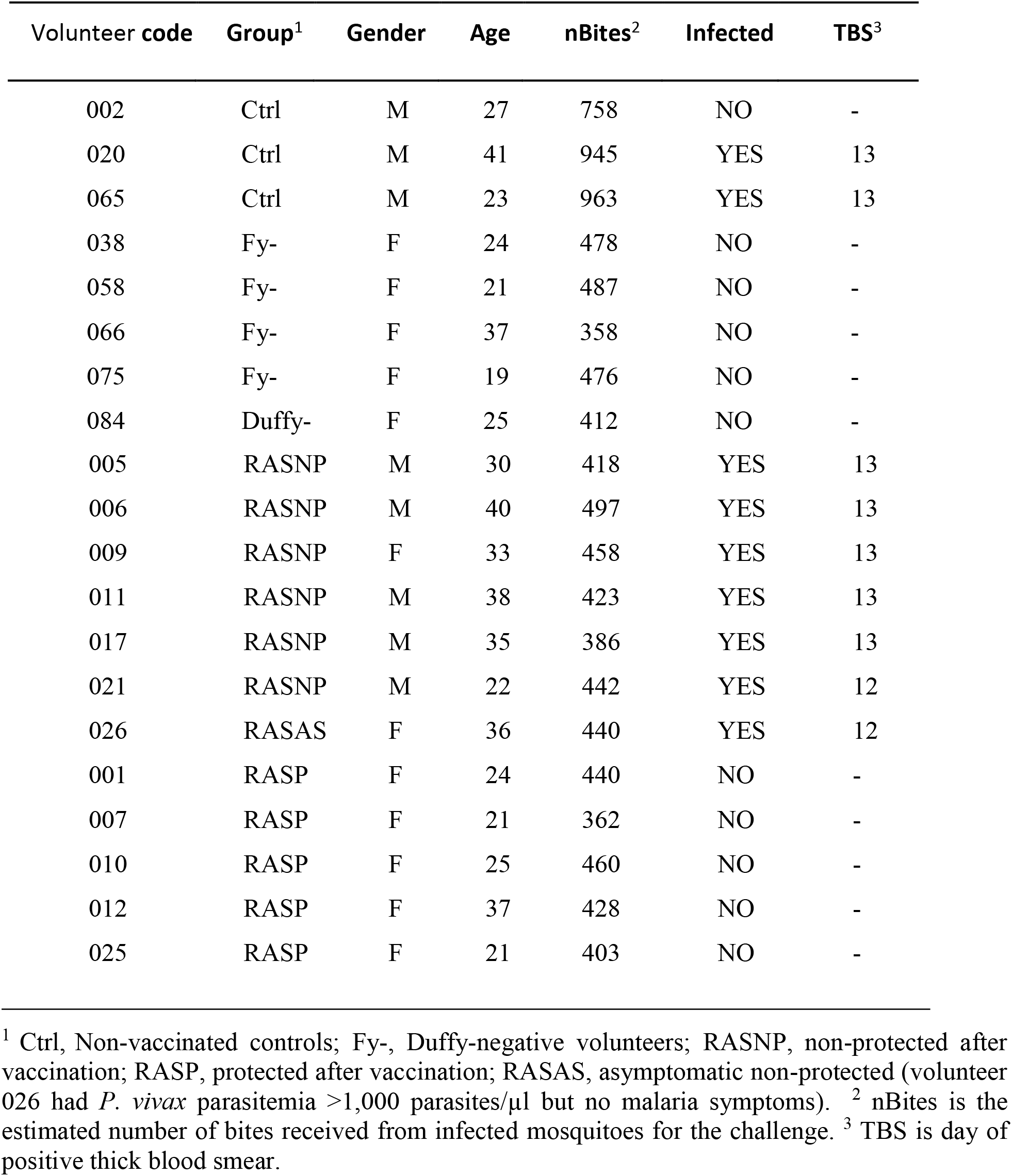
Demographics of the 20 volunteers included in the clinical trial.

Principal variance component analysis indicates that 11.6 % of the variation is between the three DPS classes of individual (Duffy-negative, protected, susceptible), 12.7 % between the three time points, and 3.3% attributable to an interaction effect between DPS status and time point. Unusually for peripheral blood profiles, gender also shows a large effect (14.4%), but this is mostly due to the association of gender with protection in the post-challenge samples as described below. Age was not a significant source of variation. Post-hoc analysis indicated that the Diagnosis day samples tend to be the most divergent, and that the susceptible (Fy+) individuals are the most distinct at the Diagnosis day, accounting for the interaction effect. Indeed, just a handful of genes differed between Baseline and Pre-Challenge at p<0.0001. Consequently, we first focused on the question of whether live parasite challenge induces different immune profiles according to clinical outcome at diagnosis day in infected individuals.

Unavoidably, the comparison of Fy- status with Fy+ is confounded with genome-wide ancestry proportions, since absence of the Fy antigen is characteristic of African populations in which natural selection at the locus has shaped protection against *P. vivax* malaria. Fig 2 confirms this genetic differentiation, showing the admixture proportions inferred using ADMIXTURE software [26] with k=3 populations, utilizing common genotypes detected by the GATK HaplotypeCaller [27] in all three samples of each individual. Red, blue, and olive colored circles at the corners of the triangle are representative samples from the 1000 Genomes Project and green circles are study participants, who tend to have higher African ancestry than the Medellin Colombians from the 1000 Genomes Project represented by the purple circles. Notably, the five outliers with greater than 40% African ancestry are the Fy- individuals.

**Figure 2.**
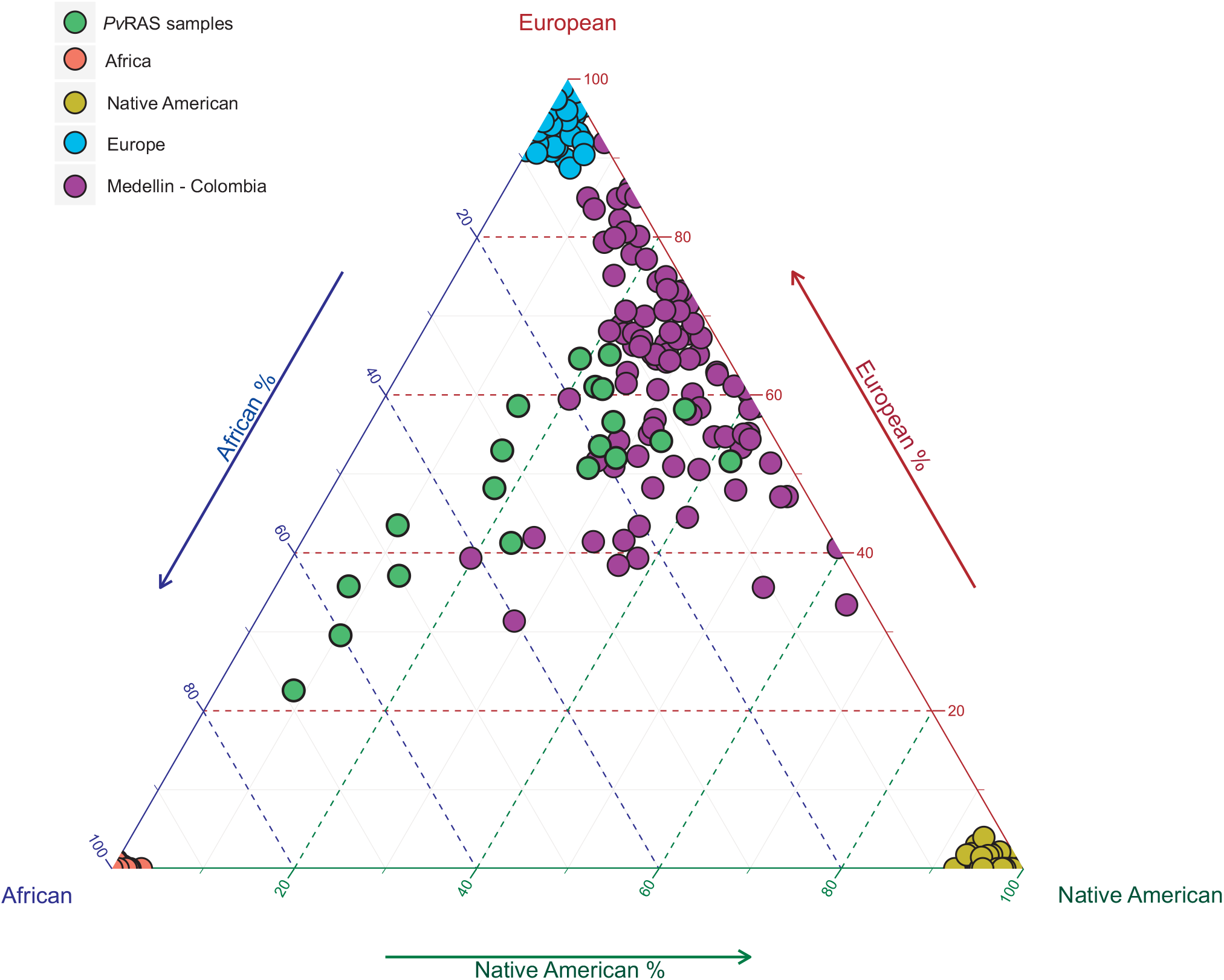
Ancestry proportions of the study participants. The three axes indicate the proportion of ancestry in each individual inferred to derive from African (left), European (right) or Amerindian (East Asian: baseline) ancestry. Ancestry proportions were inferred using ADMIXTURE [26] with k=3 populations, based on common genotypes called with the GATK HaplotypeCaller in all three samples of an individual [27], merged with the vcf-merge utility in VCF Tools [48], and matched to the 1000 Genomes Project [49] variants with PLINK [50]. Red, blue, and olive colored circles at the corners of the triangle are representative samples from the 1000 Genomes Project and green circles are study participants, who tend to have higher African ancestry than the Medellin Colombians from the 1000 Genomes Project represented by the purple circles, particularly the 5 outliers with greater than 40% African ancestry, who are Duffy negative.

### Differential Gene Expression at Diagnosis Associates with Protection or Susceptibility

Blood Transcript Module (BTM) [25,28] and Blood Informative Transcript (BIT) [29] axes are parallel methods for characterizing patterns of differential gene expression of conserved gene co-expression sets that report immune functions. The former represents coordinated gene expression as over 250 modules each annotated with respect to some aspect of immune physiology, whereas the latter collapses modules into eight larger axes of covariation. Two-way hierarchical clustering analysis of PC1 of the BTMs and BIT axis data reveals 12 clusters of co-expressed modules (8 of which include one BIT Axis), and three distinct clusters of individual (Fig 3). These three clusters largely distinguished the clinical groups: protected RAS volunteers (blue labels) separate from RAS volunteers who were susceptible (red), while the Fy- volunteers (green) form a distinct cluster more similar to the protected RAS volunteers. Two RAS non-protected individuals had aberrant profiles: one (RASNP 005) had a unique profile dominated by signatures of mitosis (multiple BTM and cell division Axis 10) indicating that the infection has temporarily induced abnormal cell cycle regulation; the other (RASNP 021) is almost identical at Diagnosis and Pre-Challenge and may be a sample mix-up.

**Figure 3.**
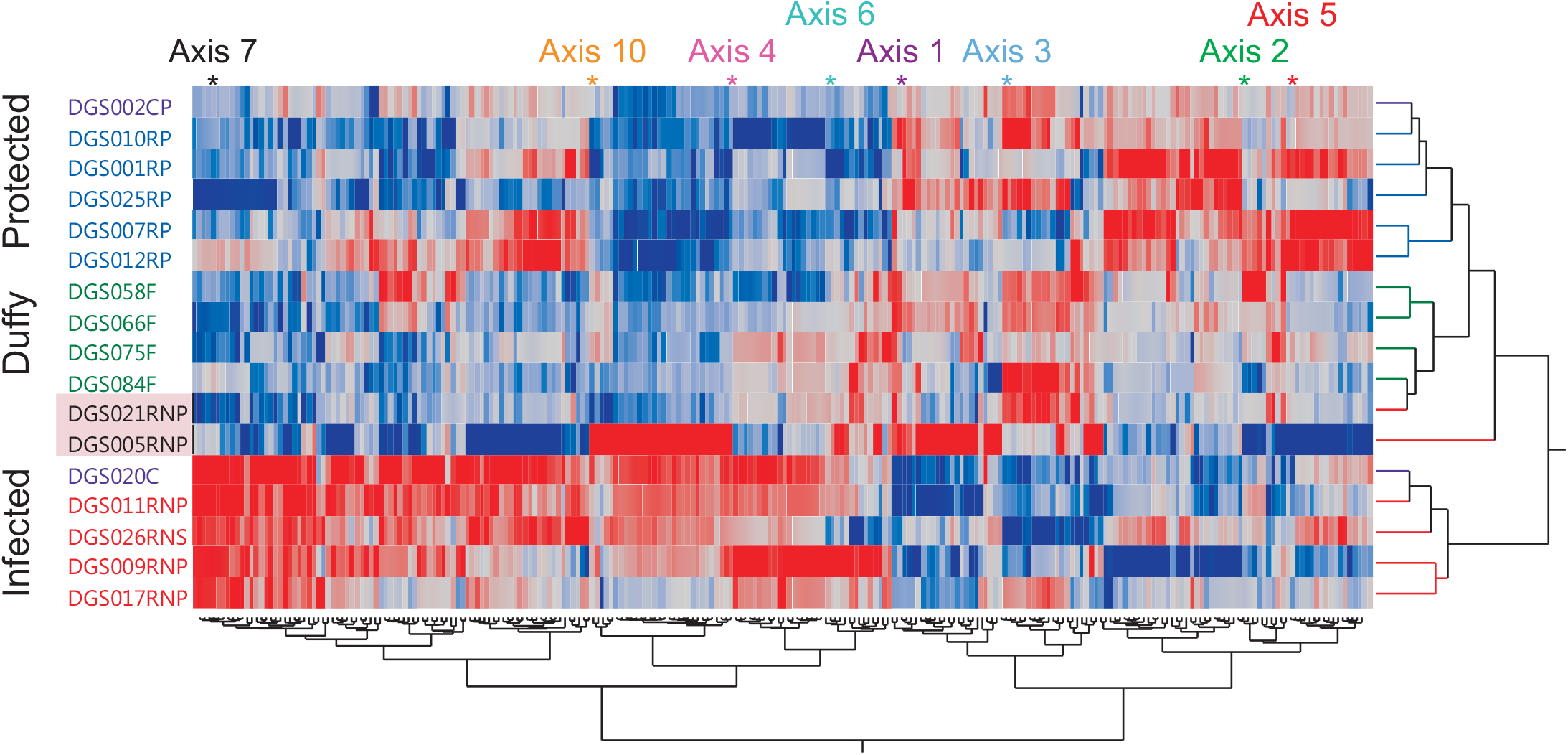
Hierarchical clustering of Blood Transcript Modules (BTMs) and Axes of Variation. The heat map shows two-way hierarchical clustering of the PC1 (columns) in each sample (rows) of 250 BTMs and 8 sets of BITs at the day of the diagnosis. Red indicates high expression, blue low, and gray intermediate, of genes in each module. Green sample labels represent Duffy negative volunteers, red labels represent susceptible RAS immunized volunteers who became infected during the challenge, blue represent protected RAS immunized volunteers, and purple, control volunteers. Nomenclature at the end of the labels means: F, Duffy negative; RNP, RAS non-protected; RP, RAS protected; C, control; CP, control protected and NS, Non symptoms.

Most apparent was the strong up-regulation of interferon response and mitosis in the non-protected RAS volunteers, contrasting with up-regulation of T-cell and B-cell signaling evident in both the Fy- and RAS protected volunteers, to varying degrees. It is particularly noteworthy that volunteer 026, who became positive for the parasite but was asymptomatic, exhibited a combination of characteristics of non-protected (interferon and mitosis response) and protected individuals (platelet and inflammation). It was also apparent that the two controls clustered with their respective clinical groups, namely 002 who did not become infected (but appears to have been exposed since his profile is related to that of the RAS protected volunteers and different from his Pre-challenge sample, not shown), and 020 who had a classical malaria profile.

A more comprehensive picture of the changes in gene expression as parasites become patent is provided by a summary of the significantly differentially expressed BTM and BITs across the 9 categories, baseline, pre-challenge and diagnosis, each subdivided into Fy- and Fy+ protected and susceptible, in S2 Table. Results for the BIT axes are summarized in S1 Fig, which confirms that T-cell signaling (Axis 1) is already induced at pre-challenge time in Fy- individuals but delayed until the diagnosis time in the RAS protected, while it does not become induced in the susceptible individuals at all. Neutrophil activity (Axis 5) is generically reduced in the Fy- even at baseline, and did not change across time within each type, with the exception of a surprising slight reduction in some of the susceptible individuals (consistent with previous report of the response of non-vaccinated naïve volunteers [16]). Interferon signaling (Axis 7) increases in most of the susceptible individuals at the time of the diagnosis, but remains unchanged in the refractory Fy- and the protected Fy+ individuals. Similarly, mitotic activity (Axis 10) is dramatically up-regulated only at diagnosis in the susceptible, non-protected vaccinees and control.

Analysis of variance of the PC1 scores of the BTM modules at day of diagnosis across the DPS categories identified 48 modules that are dysregulated with elevation associated with protection (absence of parasite) (S2a Table), and 49 modules that are associated with clinical susceptibility, being elevated in the presence of parasites (S2b Table). Examples of these modules are represented in Fig 4, which also implies that there are some differences in which modules are related to parasitemia and symptoms. The red individual (026) in the “protected” type is the asymptomatic but infected. Her module score is almost always closer to that of the sick individuals, being more protected-like for some modules (T-cell differentiation/Axis 1, cytoskeletal remodeling) but susceptible-like for others (antigen presentation, CD1 and dendritic cell receptors). The bottom row, panel C, shows three modules which are up-regulated in malarial volunteers as well as 026, involving chemokine receptors, cell division, and interferon/Axis 7 signaling, implicating these processes more in sickness or recovery than induction of sterile immunity.

**Figure 4.**
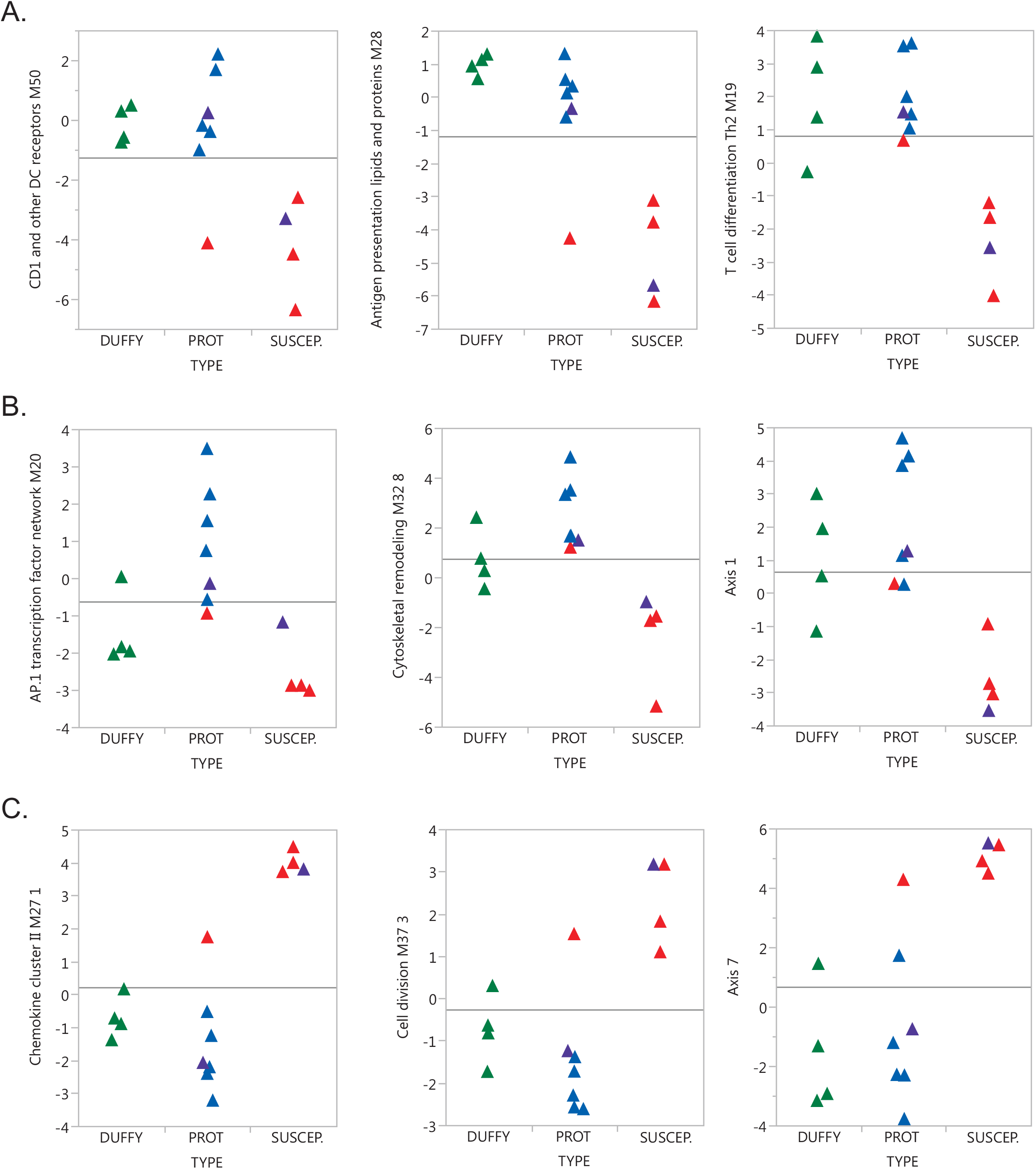
BTM and Axes distinguishing Fy^-^, protected and susceptible volunteers at diagnosis. Each plot is an example of a module or axis of variation, which differentiates volunteers who exhibited symptoms from volunteers who did not become infected. (A) 3 BTM down-regulated in susceptible individuals. (B) 2 BTM and Axis 1 up-regulated more in RAS than Fy- protected individuals. (C) 2 BTM and Axis 7 up-regulated in susceptible individuals. Green dots represent Duffy negative volunteers, red represent non-protected RAS sick volunteers, blue represent RAS protected volunteers. The red triangle in the protected RAS column represents volunteer 026 who was asymptomatic but positive for parasite, while the purple dots are the controls, one of whom was unexpectedly protected.

### Differential Expression Associated with Fy- status

The contrast of Pre-Challenge with Baseline in the Fy- individuals revealed substantial differential expression that was qualitatively similar to the comparison of Diagnosis day with Baseline for these individuals. This is as expected, since the sporozoite challenge is essentially the same as their exposure to live parasites during vaccination. Effectively, the blood gene expression is reporting a response to the presence of liver trophozoites, as RAS are unable to progress to the blood stage. To establish this further, Fig 5A plots differential expression between Fy- pre-challenge and Baseline, contrasted with differential expression between individuals protected by RAS and Baseline. All of the genes significantly differentially expressed at Diagnosis day in the RAS individuals, even though not significant at Pre-challenge in the Fy^-^, are trending in the same direction. This is particularly notable for the T-cell genes (red), but is also true of the neutrophil genes (blue) which are generally lower in Fy- individuals naturally refractory (Fig 5B), but this may also simply reflect a greater proportion of African ancestry in these individuals.

**Figure 5.**
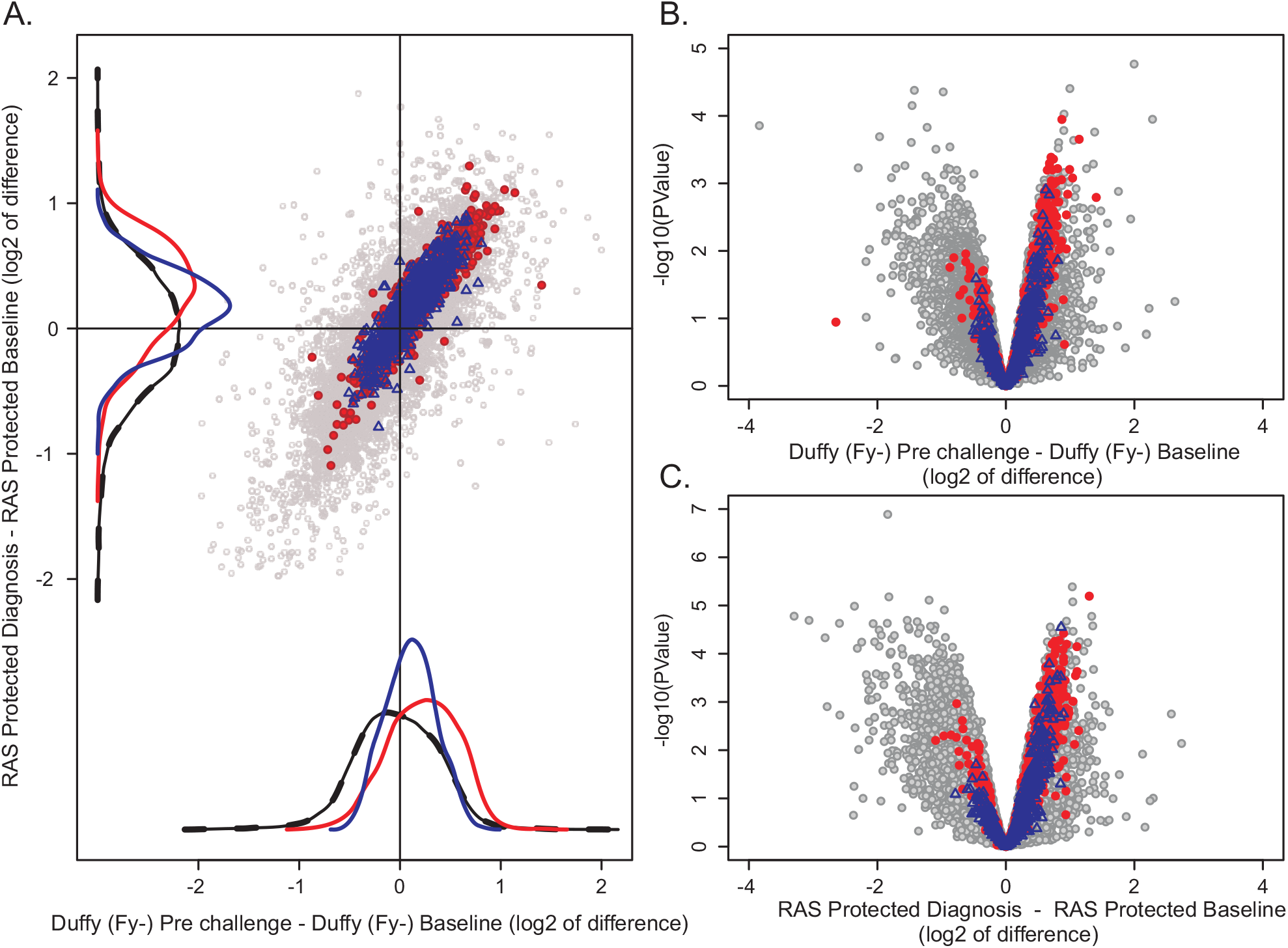
Similarities between natural and vaccine-induced protection. The scatter plot (A) shows differential expression between protected RAS volunteers post-challenge and at baseline on the y-axis (up-regulation after challenge is to the top), and between Fy^-^ volunteers pre-challenge and at baseline on the x-axis (up-regulation after vaccination to the right). Red circles represent 421 genes positively correlated with Axis 1 (r>0.5), blue circles represent 765 genes positively correlated with Axis 5 (r>0.5), and gray circles all remaining genes. Volcano plots [B,C] show the relationship between significance and difference in gene expression with significance as the negative logarithm of the p-value on the y-axis, and difference in log2 units on the x-axis with higher expression at baseline to the left.

Despite this similarity between the responses of individuals protected against live parasite, there are some very clear differences between the Fy- and Fy+ samples prior to the live sporozoite challenge: 79 genes were significantly up-regulated at FDR 5% in Fy^-^, and 658 genes are down-regulated. This result confirms that Fy status, through its influence on the *P. vivax* parasite cycle has a marked impact on the overall gene expression profile. It is likely that a different antigen repertoire is presented to the immune system in the Fy+ volunteers as consequence of the limited development of RAS as opposed to the full sporozoite development in the Fy- individuals. Fig 6 shows five examples of BTM that are differentially expressed in Fy- volunteers at pre-challenge. These include elevated gene expression related to subsets of chemokine signaling, B cell development, and integrin-mediated cell signaling, and down-regulation of extracellular matrix and complement activation (suggesting reduced general inflammation at baseline). S3 Table provides a full listing of Duffy genotype-biased gene expression modules.

**Figure 6.**
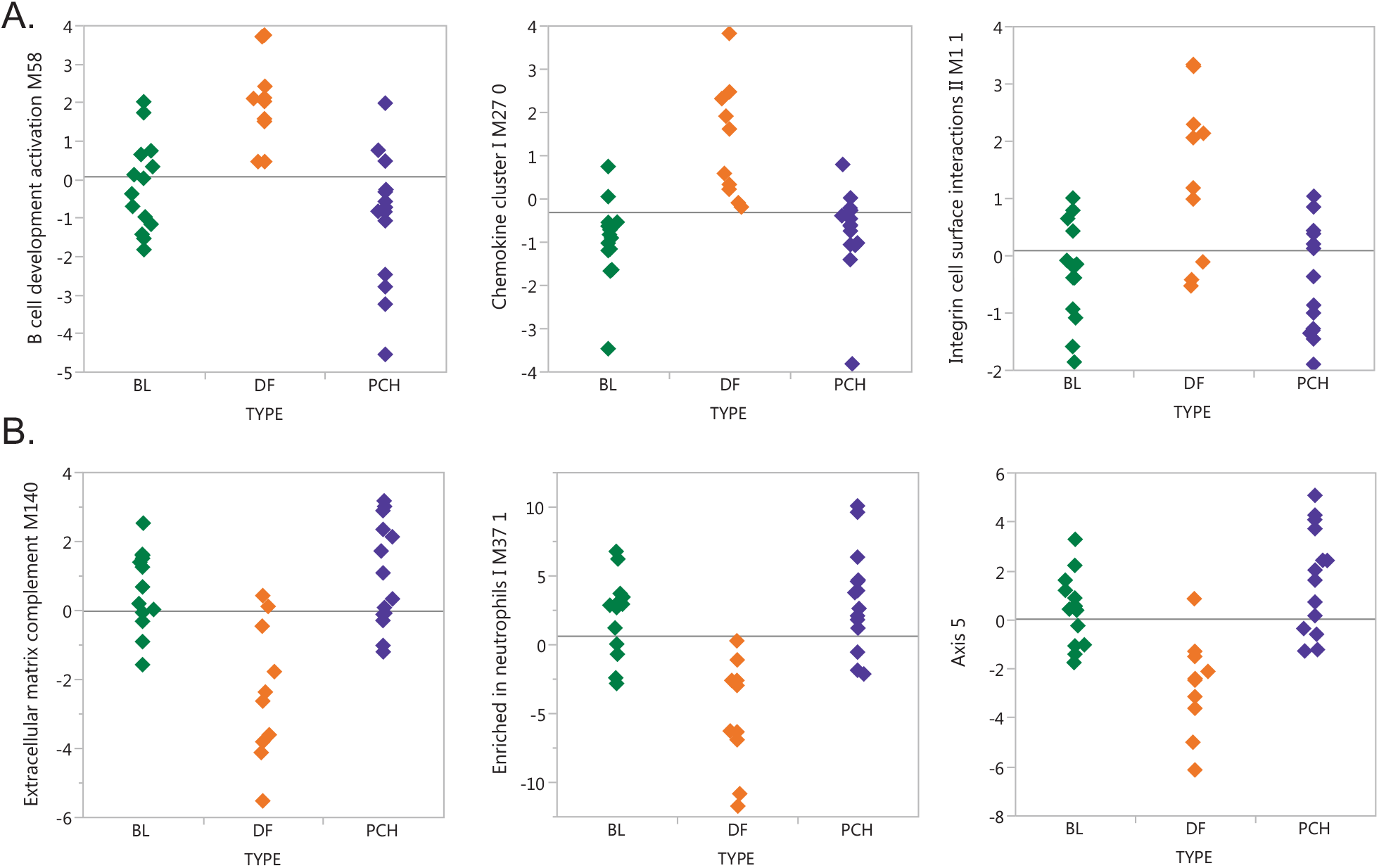
BTMs and axes that distinguish Duffy negative and positive before malaria challenge. Each plot represents a BTM or axis that differentiates Duffy negative volunteers from the other volunteers at Baseline or Pre-challenge. Green symbols represent baseline, purple pre-challenge and orange Duffy negative volunteers at either time point.

## DISCUSSION

Comprehensive BTM analysis of the immune profiles elicited by immunization with RAS *vs* live sporozoites confirms that both the B- and T-cell arms of the adaptive immune response are engaged in vaccine-induced protection, as previously shown for natural immunity (16). It is known that a B-cell response is required for regulation and elimination of infected red blood cells during the blood stage of infection [30], and correspondingly it is remarkable that 3 of 4 Fy+ infected individuals showed no sign of B-cell activation at diagnosis time as captured by Axis 3 and related BTM, while it was up-regulated after challenge in Fy+ protected individuals, indicating that as expected, B-cell activation is essential for protection.

We attempted to evaluate whether specific sub-types of B-cells may be engaged by performing FACS sorting of naïve, activated, atypical and classical B-cells. These proportions varied at diagnosis according to infection status (Fig 7A), but these differences could not be confirmed by transcriptional biomarkers of the sub-types defined in [31]. In whole blood, there is covariance of the 27 biomarker genes (Fig 7B) such that their first three principal components vary by time and infection (Fig 7C), but the covariance does not correspond to the patterns of the B-cell sub-types observed by flow cytometry analysis, so is presumably due to expressed in other immune cell types. Alternatively, a role for plasma cells is implicated by strong up-regulation of CD38 (Fig S1) and hundreds of co-expressed genes that distinguish short- and long-lived plasma cells [32,33] in the non-protected individuals (Fig S2). This suggests that they are specifically engaged in immunoglobulin-based clearance of the parasite, but sub-type specific RNA-Seq would be required to confirm this hypothesis.

**Figure 7.**
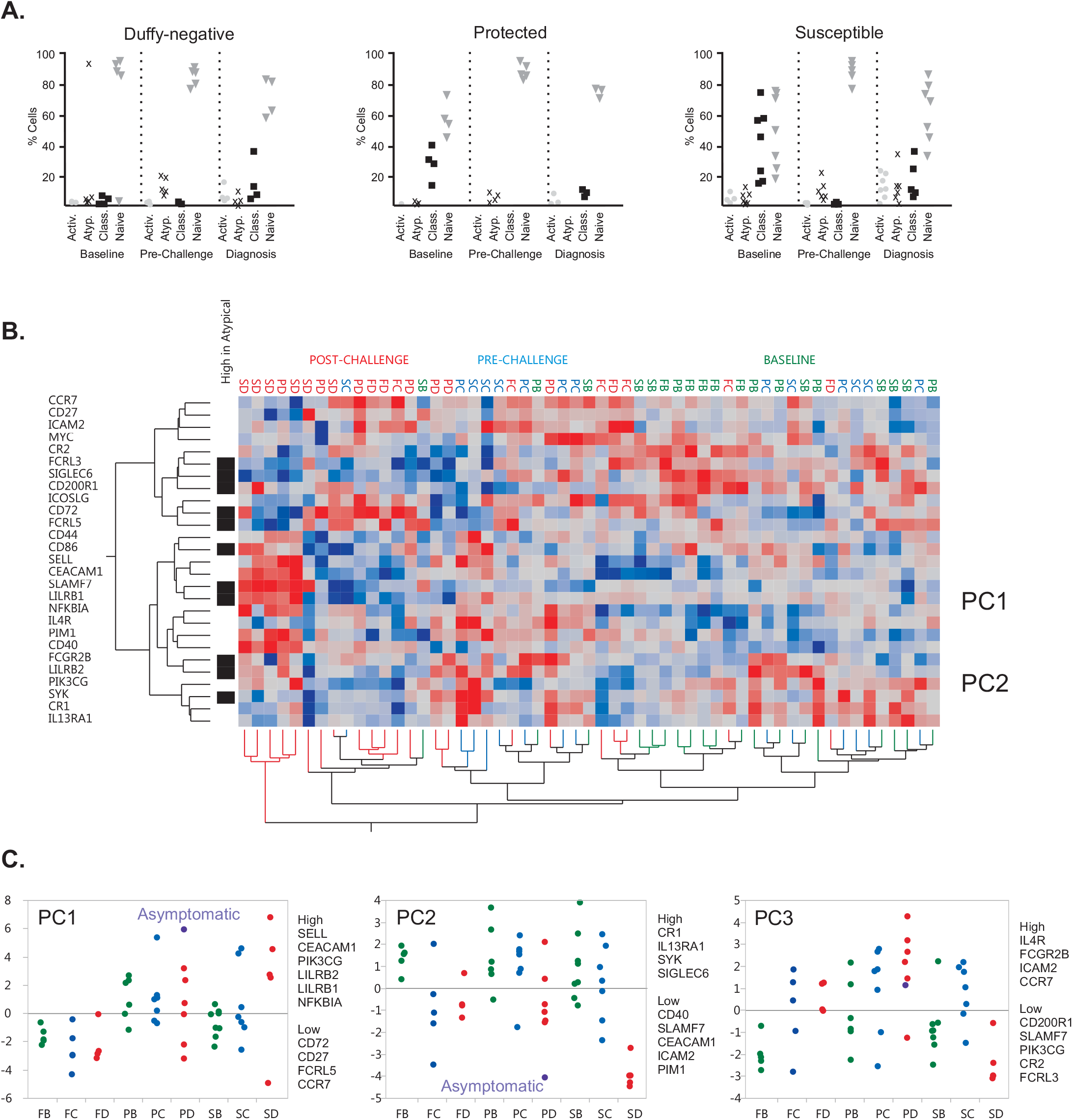
Transcriptional biomarkers of B-cell subsets do not resolve classical and atypical B cell responses. (A) Differential prevalence of activated, classical, and atypical B cell subtypes between protected and susceptible volunteers assessed by flow cytometry at Baseline and Diagnosis. (B) Heat map of gene expression of transcripts shown to distinguish atypical (black bars to left of map) from classical (remainder) B-cells [31] does not correspond with co-regulation across the three-points. Consequently, the transcript profiles cannot be used to infer abundance of the cell types detected by flow cytometry. (C) The first three principal components of this gene set identify B-cell enriched genes that are up-regulated in susceptible individuals (PC1), down-regulated in susceptible individuals (and to some extent in challenged Fy^-^) (PC2), or up-regulated in protected individuals (PC3).

T cell activation and differentiation were also up-regulated only in protected individuals, supporting previous findings that T cells proliferation is associated with *P. falciparum* malaria protection in RTS,S-immunized volunteers [34,35], including enhanced activation of memory T cells [36]. Other studies have also reported an increase in T cell responses and a delayed pre-patent period after malaria challenge [37,38], although there is some heterogeneity in lymphocyte activation levels. This may be due to a delay in the timing rather than a dampening of the lymphocyte response, since 007 and 012 volunteers, with lesser B- and T-cell gene activity also had specifically elevated activity of dozens of interferon response-related modules. In contrast to the protected volunteers, the non-protected displayed up-regulation in modules related to type I interferon response, pro-inflammatory cytokines, complement activation, cell migration, dendritic cells, and an extensive set of mitosis related modules, suggesting that volunteers who could not arrest the liver infection activated more robust immune and inflammatory responses in presence of parasite blood stages.

Previous studies have shown release of pro-inflammatory mediators like TNF and IFN-γ during malaria infections, potentially contributing to organ damage [39]. We postulate that this does not occur in protected individuals because effective vaccination facilitates an immediate recall of memory cells which either rapidly clears or arrest the liver trophozoites. Volunteer 026, who displayed parasitemia but was asymptomatic, had a hybrid profile consisting of a strong interferon response and inflammatory activation, also with differences in modules related to the cell cycle as observed previously in semi-immune individuals in response to a *P. vivax* CHMI [16]. She also had a signature of platelet activation which was recently proposed to have protective properties against malaria infection by binding infected erythrocytes and killing the parasite specifically by factor 4 activity [40,41]. Larger studies should thus enable distinction between pathways that provide vaccine-mediated sterile protection, and the ones that are more involved in malaria symptomology. In addition, many immune events that occur locally in the liver would escape detection by transcriptome profiling of peripheral blood cells. Notably, liver cells infected *in vitro* with irradiated and non-irradiated *P. falciparum* display differential gene expression as a function of sporozoite radiation status [42].

By demonstrating differential systemic responses that correlate with clinical outcomes, our study further supports the use of systems biology approaches to understand vaccine efficacy. Follow-up experimental approaches might utilize *ex vivo* parasite challenges to evaluate specific cellular contributions to immunity and protection, with a particular focus on the reasons for the female-bias, which has also been seen in other settings [43,44]. Further, profiling of sorted cell types during vaccination and in the pre-patent period days after initial infection should also reveal critical mechanisms of immunity.

## MATERIALS AND METHODS

### Clinical Study

This study is based on a single-blind clinical trial registered at ClinicalTrials.gov as NCT01082341 conducted in Colombia and described in [17], which included healthy adult volunteers, males and non-pregnant females, between 19-41 years of age, who satisfied inclusion/exclusion criteria as determined by clinical history and blood tests. Volunteers were broadly informed about the risks of participation and required to pass an oral or written exam related to the trial and its risks. The experimental protocol was approved by the Institutional Review Boards (IRB) at the Centro International de Vacunas and Centro Médico Imbanaco in Cali (Colombia), and subsequently the Georgia Institute of Technology IRB gave approval for the genomic profiling.

Fifteen malaria-naïve Fy^+^ (12 RAS and 3 mock-controls) and 5 Fy^-^ volunteers from Cali were included. The volunteers visited the clinic in Cali approximately every 8 weeks for a total of seven visits, to receive the vaccination dose, ~100 bites from *Anopheles albimanus* mosquitoes. In the case of the RAS vaccinees, mosquitoes harboring *P. vivax* mature sporozoites had been irradiated whereas the Fy- volunteers were exposed to mosquitoes harboring live non-attenuated parasites [17]. The mock-controls were immunized by exposure to the bite of non-infected mosquitoes. Two weeks after the final vaccine dose, the “pre-challenge” blood sample was drawn, and six weeks later, volunteers were challenged by exposure to the bites of 3-4 mosquitoes infected with live, non-irradiated *P. vivax* sporozoites. Twelve or thirteen days later, at first diagnosis of infection in susceptible volunteers, the final “Diagnosis” blood sample was taken, and when necessary curative anti-malarial medication was given. Two of the mock-controls contracted malaria after challenge with live parasites but one who accidentally received one round of vaccination remained uninfected. Of the 12 RAS-vaccinated individuals, six (all women) were symptom-free, one of whom nevertheless had parasites in her blood. The other six (five men, one woman) were not protected and developed malaria. As expected, all five Fy- volunteers (all women) were protected from blood infection.

### B-cell phenotype analysis

Peripheral Blood Mononuclear Cells (PBMC) were isolated by ficoll-histopaque (Sigma-Aldrich, USA) density gradient according to the manufacturer’s instructions. Briefly, fresh PBMC were incubated with mouse mAbs specific for human B cell markers conjugated to fluorophores. The monoclonal panel included: PerCP-CD19, PE-CD10, APC-Cy7-CD20, PerCy7-CD21, APC-CD27, and FITC-CD95 (BD Science, San Jose, CA). Isotype matched controls FITC-IgG (BD Pharmigen, San Jose, CA) were used in all experiments. Samples were acquired using a FACSCanto II flow cytometer with FACSDiva software (BD Biosciences, San José, USA). Data were analyzed using FlowJo software (Tree Star, Inc).

### RNAseq

Whole blood RNA was prepared from ~1 mL of blood drawn from each of the 20 individuals at three time points, namely Baseline (prefix B), Pre-Challenge (C) and at Diagnosis (D). Each sample was mixed with 2 mL of stabilizing buffer in a Tempus™ tube (Applied Biosystems, CA, USA), which preserves whole blood RNA at -80°C indefinitely. Whole blood (predominantly leukocyte) mRNA was extracted using Tempus Blood RNA Tube isolation kits following the protocol provided by Applied Biosystems. Sample RNA quality was determined based on the

Agilent Bioanalyzer RNA Integrity score (RIN: S1 Table). Two RNA samples, one from prechallenge time of Fy+ control group and the other from diagnosis time of Fy- group, were severely degraded and therefore not included for sequencing. A few samples had RIN <6, but these were not found to be outliers in the analysis.

RNAseq libraries were prepared using the Illumina TruSeq Stranded mRNA Low Throughput sample preparation kit and protocol. Short read sequencing was performed in rapid run mode with eight samples per lane on an Illumina HiSeq 2500 at the Georgia Institute of Technology, generating 100 bp single-end libraries with an average of 31.8 ± 6.1 million single end reads per sample. The RNASeq dataset has been deposited into the Gene Expression Omnibus archive (GEO) under accession number GSE85263, including short read deposition in the SRA.

### Data analysis

The quality of raw RNAseq reads in Fastq files was evaluated using FastQC software. The overall alignment rate was 93%, but three samples (B025, DGS006 and DGS065) failed QC and were excluded. The 100 bp single-end reads were aligned to the human genome (hg19/GRCh37 assembly with the UCSC reference annotation) using Bowtie via Tophat2 [45]. Transcript abundance was estimated at the level of the whole gene as counts per million-mapped-reads (cpm) values using HTSeq [46]. Scale Trimmed Mean of the M-values (TMM) normalization was then performed using edgeR from Bioconductor [47], and differential expression was assessed using the linear modeling framework for RNAseq data in edgeR, selecting transcripts significant at the Benjamini-Hochberg false discovery rate of 10% (corresponding to a p-value of ~0.0001).

All results were verified by a second normalization model of the log2 transformed data, using the SNM package in R [48] to fit individual as an adjustment variable with time point and clinical status as biological variables, facilitating analysis of the relative changes in expression after adjustment for inter-individual differences. The contributions of time point, gender and clinical status (DSP, Duffy, Susceptible or Protected) to the overall gene expression variation were estimated using the Principal Variance Component Analysis routine in the Basic Expression Workflow in JMP Genomics (SAS Institute, Cary NC). Since all differential expression results were qualitatively the same as the TMM analysis, we only report the TMM.

Ancestry proportions of the study participants were inferred using ADMIXTURE [26] with k=3 populations, based on common genotypes called with the GATK HaplotypeCaller in all three samples of an individual [27], merged with the vcfmerge utility in VCF Tools [49], and matched to the 1000 Genomes Project [50] variants with PLINK [51]. Red, blue, and olive colored circles at the corners of the triangle are representative samples from the 1000 Genomes Project and green circles are study participants. These tend to have higher African ancestry than the Medellin Colombians from the 1000 Genomes Project represented by the purple circles. The five outliers with greater than 40% African ancestry close to the left hand edge of the triangle in Fig 2 are the Fy negative individuals.

### Blood Transcription Modules (BTMs) and Blood Informative Transcripts (BIT)

Modular analysis of transcript abundance was based on the reconstruction of several hundred gene networks from integrative analysis of >30,000 transcriptomes in 500 public studies [25]. These modules are described as Blood Transcription Modules (BTMs) and disclose distinct aspects of peripheral blood gene expression, including transcriptional indications of antibody responses to vaccination and other immunological functions. In parallel, we also used Blood Informative Transcript (BIT) analysis, which focuses on 7 common axes of variation that were detected in multiple human peripheral blood gene expression datasets of healthy individuals that have been found to consistently co-vary in peripheral blood [29]. Each axis includes between one hundred and several thousand genes that gene set enrichment analysis suggests are involved in particular immune functions, broadly speaking, T cell signaling (Axis 1), reticulocyte number (Axis 2), B cell signaling (Axis 3), inflammation/neutrophil signaling (Axis 5) and Interferon signaling (Axis 7). Axis 4 appears to be related to house-keeping, Axis 6 is as yet undefined, while a newly identified Axis 10 first reported here is enriched for regulation of mitosis and cell division. These axes are related to 28 modules of co-expressed genes described in [28].

Both BTM and BIT Axes are computed as first principal components (PC1) of the co-regulated genes, either all of the genes in each module, or the 10 axis-defining BIT for each Axis (almost identical scores are obtained with 5 or 20 transcripts [29]). These scores were contrasted with respect to the clinical status groups across the three time points, using standard parametric t-tests or analysis of variance. Two-way hierarchical clustering of the BTM and BIT scores was used to assess the overall functional relatedness of samples and modules, using Ward’s method implemented in JMP Genomics. Analyses of the correlation of specific gene sets among groups were performed with custom scripts in R, accessing datasets referenced in the text.

## ACKNOWLEDGEMENTS

We particularly appreciate the assistance of Dalia Arafat-Gulick with study management in Atlanta; in Cali, the entire team and especially the volunteers for their contributions to the irradiated sporozoite vaccine clinical trial; Eliana Ortiz and Angélica Castellanos for flow cytometry technical processing at Caucaseco Research Center; and Rabin Tirouvanziam and Frances Eun-Hyung Lee at Emory University for discussions.

## SUPPORTING INFORMATION CAPTIONS

**S1 Figure.**
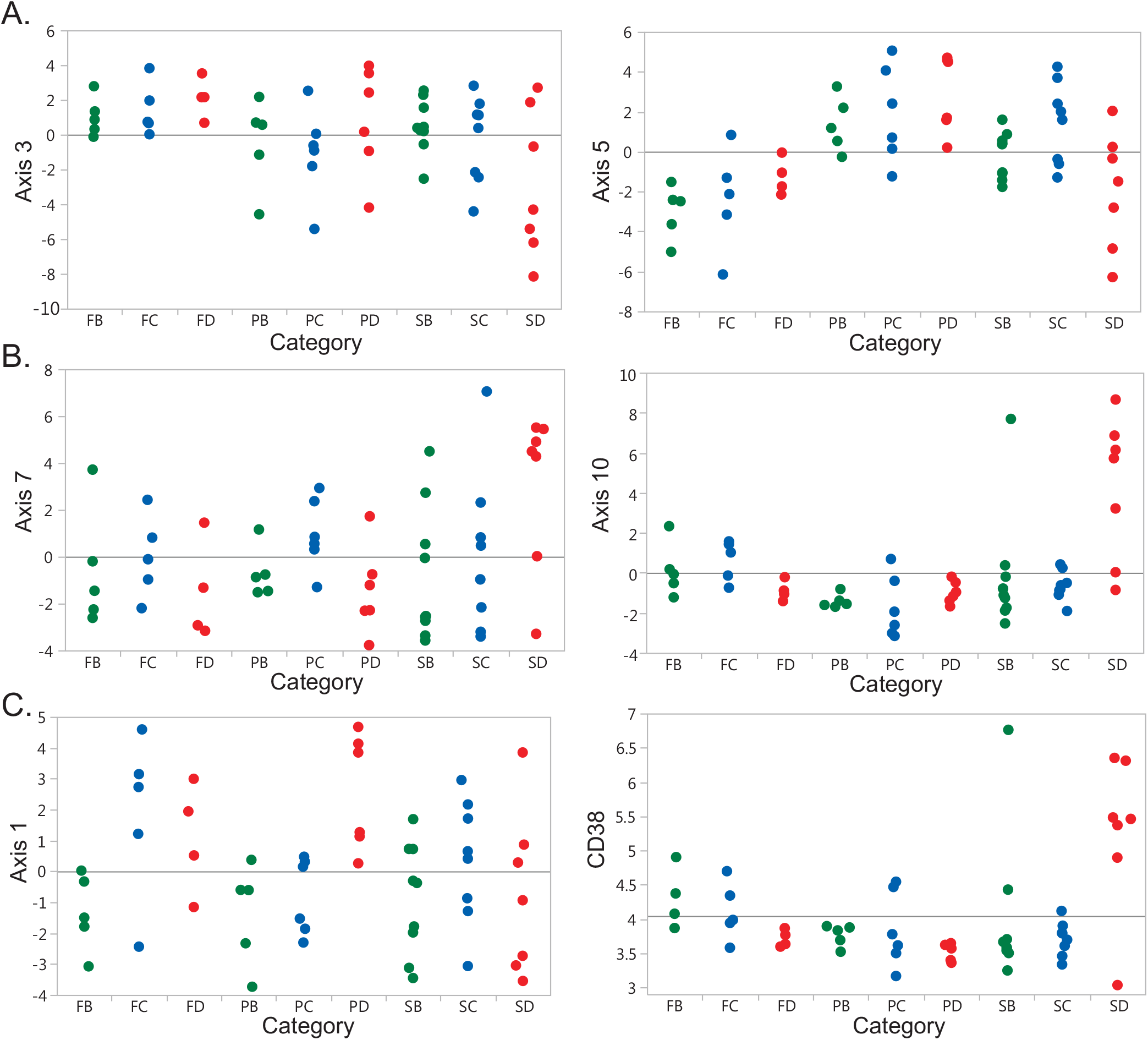
Differential expression of axes of variation. Each panel shows the PC1 score for the indicated Axis for individuals at Baseline (green, B), Pre-Challenge (blue, C) or Diagnosis Day (red, D), with from left to right Duffy negative (left, F), Protected (middle, P), or Susceptible (right, S). Panels are top to bottom, left to right: T-cell signaling (Axis 1), B-cell signaling (Axis 3), Neutrophil signaling (Axis 5), Interferon response (Axis 7), Mitosis (Axis 10), and expression of the CD38 gene.

**S2 Figure.**
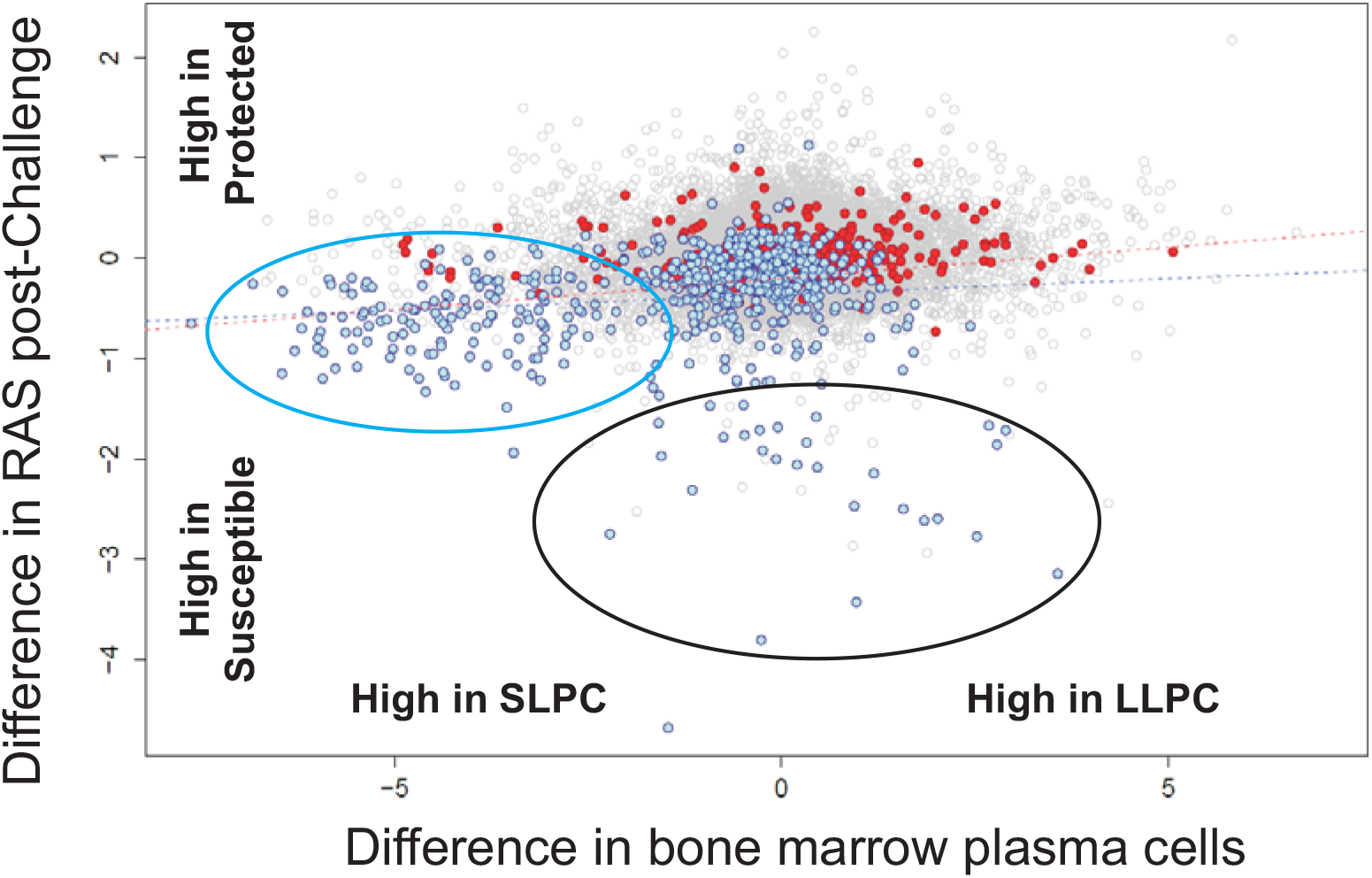
Apparent up-regulation of Short-Lived Plasma Cells in susceptible individuals. The x-y plot shows all genes correlated with CD38 in this study at r>0.5, with 607 genes positively correlated in blue and 323 negatively in red. The x-axis compares differential expression between the type A “SLPC” and type D “LLPC” cells from the bone marrow of three human donors [34], and the y-axis shows the difference between the susceptible and protected RAS individuals post-challenge. The regression for all colored genes is 0.24, *P*<10^−12^. Almost all genes >2-fold higher in SLPC are also up-regulated in the susceptible individuals, but the two dozen genes with the highest expression in susceptible individuals do not show a consistent pattern of differentiation in the plasma cells.

